# Development of a simple and accurate molecular tool for *Spodoptera frugiperda* species identification using LAMP

**DOI:** 10.1101/2020.04.07.029678

**Authors:** Juil Kim, Hwa Yeun Nam, Min Kwon, Hyun Ju Kim, Hwi-Jong Yi, Sabine Haenniger, Melanie Unbehend, David G. Heckel

**Affiliations:** Highland Agriculture Research Institute, National Institute of Crop Science, RDA, 25342 Pyeongchang, Republic of Korea; Crop foundation Division, National Institute of Crop Science, RDA, 55365 Wanju, Republic of Korea; Crop Production Technology Research Division, National Institute of Crop Science, RDA, 50424 Miryang, Republic of Korea; Department of Entomology, Max Planck Institute for Chemical Ecology, 07745 Jena, Germany

**Keywords:** *Spodoptera frugiperda*, fall armyworm, mitochondrial genome, LAMP, diagnostic PCR

## Abstract

The fall armyworm, *Spodoptera frugiperda* is a native species in the Americas. However, nowadays it is one of the most serious invasive lepidopteran pests in African and Asian countries. *S. frugiperda* has been spread very quickly after the first outbreak was reported in many countries. Based on mt genome sequence alignment, *S. frugiperda* specific sequence region was identified in tRNAs coding region between NADH dehydrogenase, ND3 and ND5. By using this unique region, species diagnostic primers were designed and applied in LAMP (lamp loop mediated isothermal amplification) assay as well as conventional PCR to identify the field-collected samples of *S. frugiperda*. Optimal incubation condition of LAMP assay was 61°C for 90 minutes with 4 LAMP primers, and additional loop primer increased the amplification efficiency. Also, wide range of DNA concentration responded in LAMP assay and minimum detectable DNA concentration was 10 pg. This LAMP assay was also applied in DNA releasing technique from larval and adult sample, without DNA extraction, 95°C incubation for five minutes of the tissue sample. This new molecular diagnostic method is easy to use and accurate. It possibly applied in intensive field monitoring of *S. frugiperda* and its ecological studies.

## 1. INTRODUCTION

The fall armyworm, *Spodoptera frugiperda* (J.E. Smith) (Lepidoptera: Noctuidae), is a polyphagous migratory pest that feeds on over 80 species of plants, mostly known as a major pest of maize and various cereals (Montezano et al., 2018; Spark, 1979). There are two morphologically indistinguishable host-plant specific strains of *S. frugiperda*, the “corn strain” and the “rice strain”. The corn strain prefers to feed on corn, cotton, and sorghum, while the rice strain prefers rice and various pasture grass (Dumas et al., 2015). These two strains have been hypothesized to start diverging about 2 Myr ago and continue to diversify (Gouin et al., 2017).

The females lay eggs on abaxial leaf surfaces of host leaves and lay up to 200 eggs in an egg mass on a single host (Kumela et al., 2019). At high densities, *S. frugiperda* larvae can completely defoliate their host plants and can cause severe yield losses in many economically important crops. In Brazil, attacks from *S. frugiperda* can reduce the corn grain yield up to 34%, causing losses of 400 million US dollars annually (Lima et al., 2010).

*S. frugiperda* is native to the tropical and subtropical Americas but was first detected in western and central Africa in early 2016 (Goergen et al., 2016) and has spread rapidly across almost all sub-Saharan countries. Subsequently, it was verified in several Asian countries including India (Sisay et al., 2018), Thailand, Myanmar, and Bangladesh in 2018, and recently China (Jing et al., 2019). In Korea, the first observation of *S. frugiperda* was made in cornfields at Jeju in early June 2019 (Seo et al., 2019). By the end of August 2019, *S. frugiperda* larvae were detected in cornfields of eight provincial regions in South Korea. In most cases, at least two or more other lepidopteran pests, such as *S. litura* (Fabricius), *S. exigua* (Hübner), *Mythimna separata* (Walker) and *Helicoverpa armigera* (Hübner), were observed in the same cornfields. Egg mass morphology, larval morphology and plant damage may be similar for different species. Therefore, for effective pest management and monitoring of invasive *S. frugiperda*, an effective identification method is required to distinguish it from similar species. Molecular diagnostics to distinguish the two host strains of *S. frugiperda* are based on conventional PCR, restriction digestion, and sequencing of part of the mitochondrial COI gene (Cock et al., 2017; Otim et al., 2018) and sequencing of part of the nuclear-encoded, sex-linked Tpi gene (Jing et al., 2019). Species identification utilizes random amplification polymorphic DNA (RAPD)-polymerase chain reaction (PCR) (Martinelli et al., 2006), amplified fragment length polymorphism (AFLP) (Clark et al., 2007; Martinelli et al., 2007) and real-time PCR based on the mitochondrial cytochrome b gene to distinguish among *S. eridania, S. frugiperda, S. littoralis*, and *S. litura* (Van de Vossenberg & Van der Straten, 2014). However, these molecular tools are expensive, time consuming, and depend on specialized equipment. A simpler technique, termed loop-mediated isothermal amplification assay (LAMP), is also widely used for the rapid identification of pest species (Blaser et al., 2018; Blaser., 2018; Hsieh et al., 2012; Choi et al., 2016). The LAMP assay was developed for rapid, simple, effective and specific amplification of DNA. It is performed under isothermal conditions that require a set of four primers, a strand-displacing DNA polymerase, and a water bath or heat block to maintain the temperature at about 65 °C following a one-time denaturation at 95 °C (Notomi et al., 2000) or one step incubation at about 65 °C (Nagamine et al., 2002).

Following the first infestation of *S. frugiperda* in Korea, there is great demand from agricultural research, extension services, and farmers for diagnostic methods for these species. Therefore, we present a method based on LAMP based on specimens collected in Korea and other sequences from GenBank. This method should be useful in assisting effective pest management of *S. frugiperda*.

## 2. MATERIALS AND METHODS

### 2.1 Sample collection and DNA extraction

Three Korean populations were collected from Jeju (33°23.18 N, 126°37.40 E), Muan (34°57.29 N, 126°20.46 E), and Miryang (35° 29.29 N, 128° 44.31 E). Two African populations originated from Nigeria and Benin and were reared in the Department of Entomology, Max Planck Institute for Chemical Ecology, Jena, Germany. Genomic DNA was extracted with DNAzol (Molecular Research Center, Cincinnati, OH) and quantified by Nanodrop (NanoDrop technologies, Wilmington, DE, USA).

### 2.2 Mitochondrial genome sequencing

For mitochondrial genome sequencing, the Miseq platform was used and more than 1 Gb were sequenced in each sample. To assemble these data, CLC Assembly Cell package (version 4.2.1) was used. After trimming raw data using CLC quality trim (ver. 4.21), assembly was accomplished using the CLC de novo assembler with dnaLCW. Assembled sequences were confirmed by BLASTZ (Schwartz et al., 2003). The GeSeq program was used for annotation (Tillich et al., 2017) and the result manually checked based on alignment of other *Spodoptera* species mitochondrial genomes using MEGA 7 (Kumar, Stecher, & Tamura, 2016). The circular mitochondrial genome map was generated by OGDraw program (Lohse et al., 2007).

### 2.3 Phylogenetic analysis and primer design

Molecular phylogenetic analysis of mitochondrion genomes was inferred by using the maximum likelihood method implemented by MEGA 7 with bootstrapping (Kumar et al., 2016; Sanderson & Wojciechowski, 2000). Mitochondrial genome sequences of other *Spodoptera* species downloaded from GenBank, NCBI, and used the previously reported sequence reference of rice and corn strain of *S. frugiperda* (Gouin et al., 2017). For comparative analysis, mitochondrial genomes were aligned using mVISTA (Frazer et al, 2004; Mayor et al., 2000). Based on the global alignment result, partial sequences were re-aligned for LAMP primer design using PrimerExplorer V5.

### 2.4 LAMP and conventional PCR

WarmStart® LAMP Kit (New England Biolabs, Ipswich, MA) used for LAMP assay. General protocol of LAMP was flowed by manufacture’s guideline in a 25 μl reaction mixture. For the conventional PCR, TOYOBO KOD-FX Taq™ (Toyobo Life Science, Osaka, Japan) used in this study. Species-specific primers with the following PCR amplification protocol: a 2-min denaturing step at 94°C and a PCR amplification cycle consisting of denaturing at 94°C for 20 s, annealing at 60°C for 20 s, and extension at 68°C for 30 s, which was repeated 35 times. The amplified DNA fragments were separated using 1.5% agarose gel electrophoresis, visualized with SYBR green (Life Technologies, Grand Island, NY). Used insect samples were collected in Korean cornfields except *S. littoralis*. MPI-CE lab strain of *S. littoralis* and mitochondrial genome sequenced *S. frugiperda* samples (Jeju, Muan and Miryang) used. Pheromone traps were used as previously reported to collect samples (*S. exigua, S. litura, Mythimna separate*, and *Helicoverpa armigera*). Traps were set in Pyeongchang (37°40.53 N, 128°43.49 E), Hongchen (37°43.35 N, 128°24.33 E) and Gangneung (37°36.56 N, 128°45.59 E) (Kim et al., 2018). DNA samples were prepared using DNAzol from trapped adults. Two or three biological DNA samples were used in each LAMP and PCR.

## 3. RESULTS

### 3.1 Mitochondrial genome sequencing and primer design

Mitochondrial genomes of the reference corn strain and reference rice strain of *S. frugiperda* were extracted from the genome publication (Gouin et al., 2017). Another entire mitochondrial genome was reported in 2016 from China (15,365 bp) (Liu et al., 2016) and in 2019 from Korea (15,388 bp) (Seo et al., 2019). We sequenced individuals from cornfields in three Korean localities: Jeju (GenBank MN599981: 15,368 bp), Muan (MN599982: 15,368 bp), and Miryang (MN599983: 15,387 bp). Additionally, two African populations, one collected in Nigeria (MN599980: 15,400 bp) and the other in Benin (MN803322: 15,369 bp) were sequenced (Table 1). Among the five *S. frugiperda* populations, 15 indels and 232 SNPs were identified (Supplementary Information). Mitochondrial genome size varied from 15368 bp to 15400 bp and the sequence of the Jeju population was identical to the Muan population. Three indels and one SNP differed between Jeju and Miryang. The GC content was 18.7 % except for Benin (18.9 %). The three Korean populations grouped with the rice reference strain and the Nigerian population, while the Benin sample grouped with the corn reference strain (Fig. 1A).

**Table 1.** Summarized NGS sequencing and assembly information of three Korean and two African *Spodoptera frugiperda* populations

| Sample name |  | Input reads | Trimmed reads |  | Coverage (x) | mt genome length (bp) |
| --- | --- | --- | --- | --- | --- | --- |
| Korea | MN599981 | 3,535,728 | 3,232,363 | 91.42% | 107.81 | 15,368 |
| Korea | MN599982 | 4,546,290 | 4,015,292 | 88.32% | 194.66 | 15,368 |
| Korea | MN599983 | 3,159,382 | 2,790,201 | 88.31% | 155.71 | 15,387 |
| Nigeria | MN599980 | 3,839,658 | 3,524,506 | 91.79% | 204.54 | 15,400 |
| Benin | MN803322 | 5,183,990 | 4,789,700 | 92.39% | 426.04 | 15,369 |

Miseq platform was used for NGS sequencing.

**Fig. 1.**
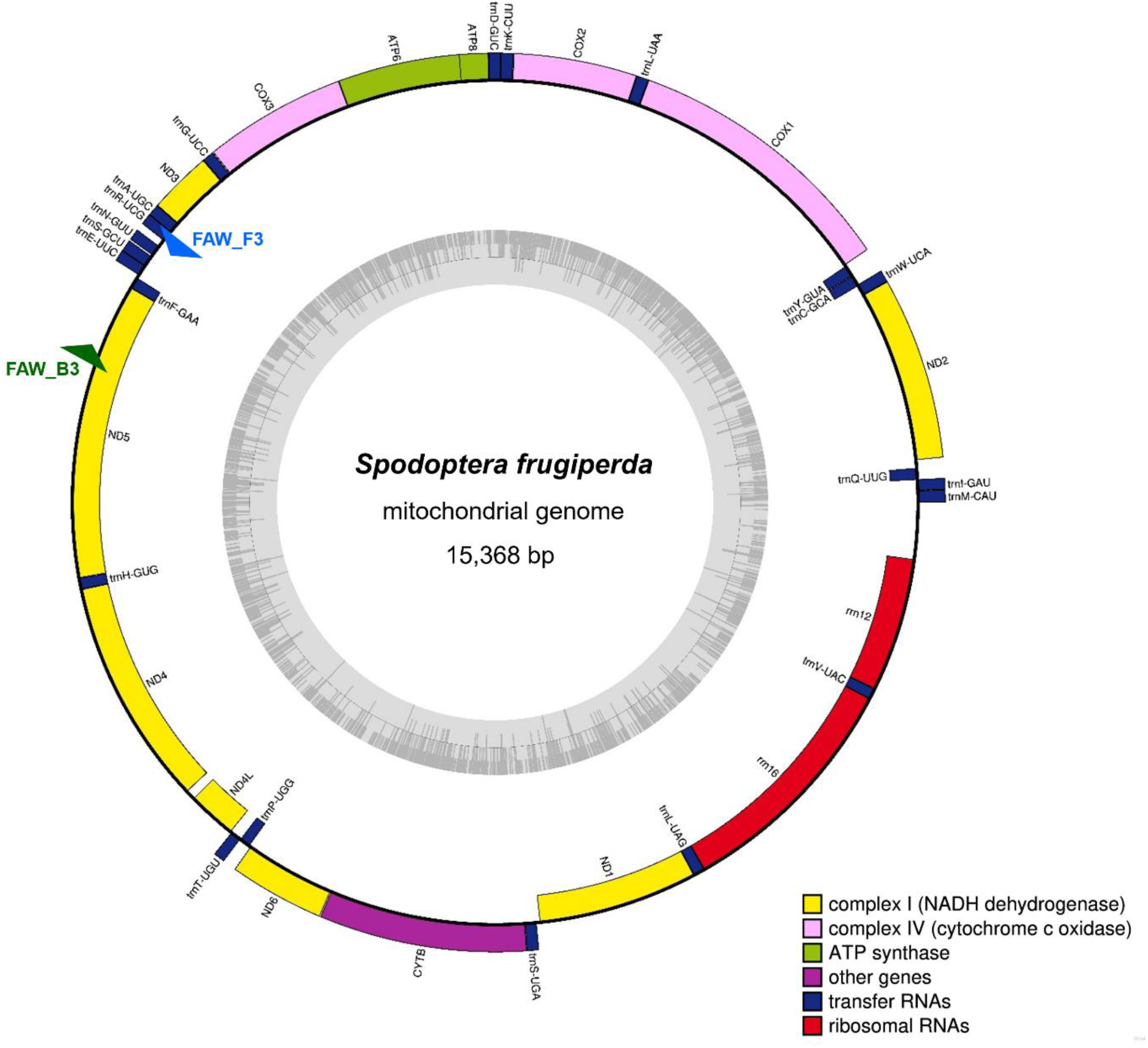
Comparison of entire mitochondrial genomes of *Spodoptera* species and strains of *S. frugiperda*. (A) Phylogeny inferred using maximum likelihood under MEGA7. (B) Schematic diagram of the genes and their flanking regions showing the sequence diversity in mVISTA. UTR, D-loop denotes untranslated region and displacement-loop, respectively.

Based on the global mVISTA alignment results, conserved regions among *Spodoptera* species and variable regions were observed (Fig. 1B). Just as for other Lepidoptera, the mitochondrial genome includes 13 protein coding genes: NADH dehydrogenase components (complex |, ND), cytochrome oxidase subunits (complex VI, COX), cytochrome oxidase b (CYPB) and two ATP synthases; two ribosomal RNA genes and 22 transfer RNAs (Fig. 2).

**Fig. 2.**
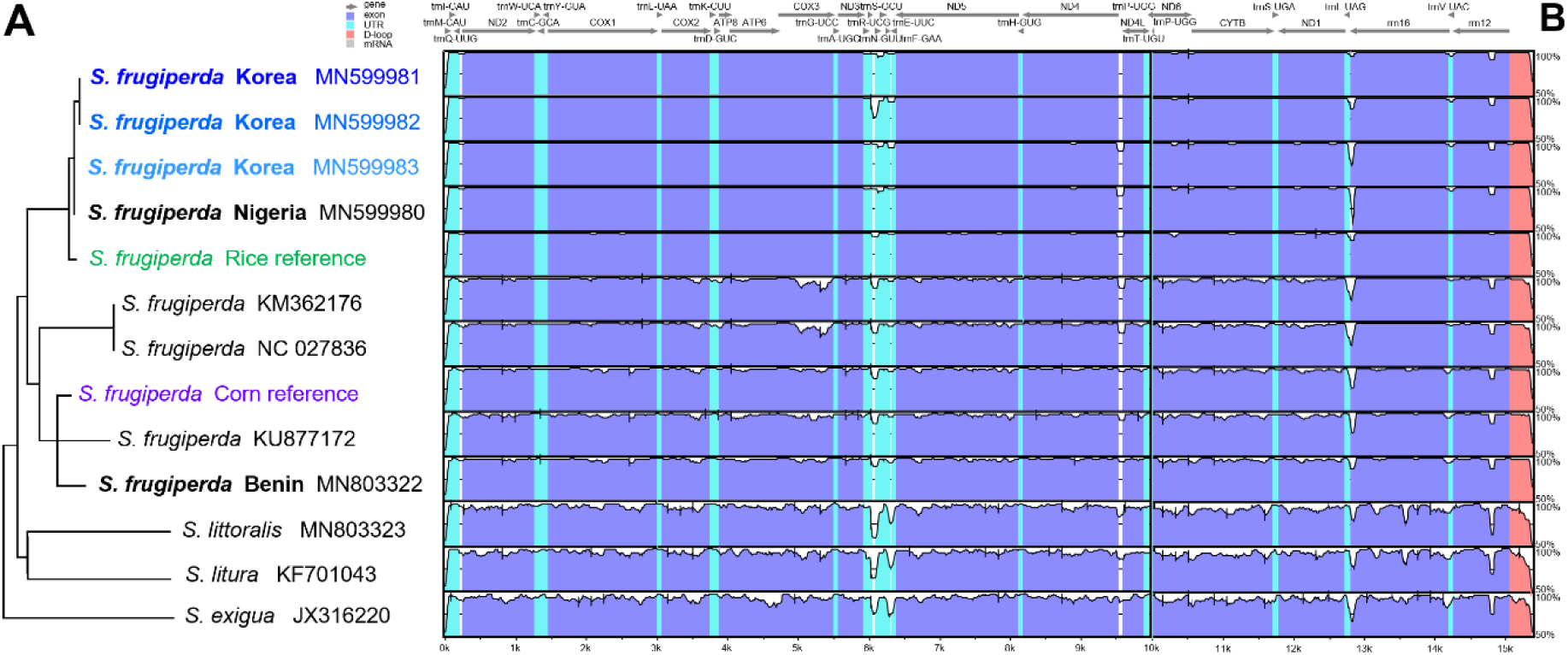
Organization of the mitochondrial genome of *Spodoptera frugiperda* from Jeju, Korea (GenBank MN599981). ND: NADH dehydrogenase components (Complex I) in yellow. COX: cytochrome oxidase subunits (Complex VI) in pink. ATP synthase in green. CYPB: cytochrome oxidase b in purple. Ribosomal RNA genes in red, tRNA genes in blue. Noncoding regions are not colored. Blue and green triangles denote species-specific primers FAW_F3 and FAW_B3, respectively.

For primer design we focused on a region including part of HD5, the tRNA-R and short non-coding region between primers FAW_F3 and FAW_R3 (Fig. 2). Primer sequences are given in Table 2 and primer-binding regions shown in Fig. 3. Primer FAW_F3 is the main diagnostic primer and along with FAW_B3 (Fig. 4A) and FAW_UR in (Fig. 4B) will amplify a product only from *S. frugiperda* mtDNA. Primer BAW_DF along with FAW_UR will amplify a product only from *S. exigua* mtDNA (Fig. 4C). These primer pairs can distinguish *S. frugiperda* and *S. exigua* by conventional PCR (Fig. 4). Primers FAW_F3 and FAW_B3 amplify the larger region that participates in the LAMP reaction, only from *S. frugiperda*. The dumbbell structure that is further amplified has one strand generated by primer BIP, which consists of B1c fused to B2 and primes from B2c on the top strand (Fig. 3). The other strand is generated by primer FIP, which has F1c fused to F2 and primes from F2c on the bottom strand. Primers FAW_F3 and FAW_B3 then further amplify the region by strand displacement through these double-stranded products. The base-paired regions in the dumbbell structure can also prime extension along with the BIP and FIP primers. Additionally, loop primers (LF and LB) could potentially contribute even more to the amplification (Nagamine et al., 2002).

**Table 2.**
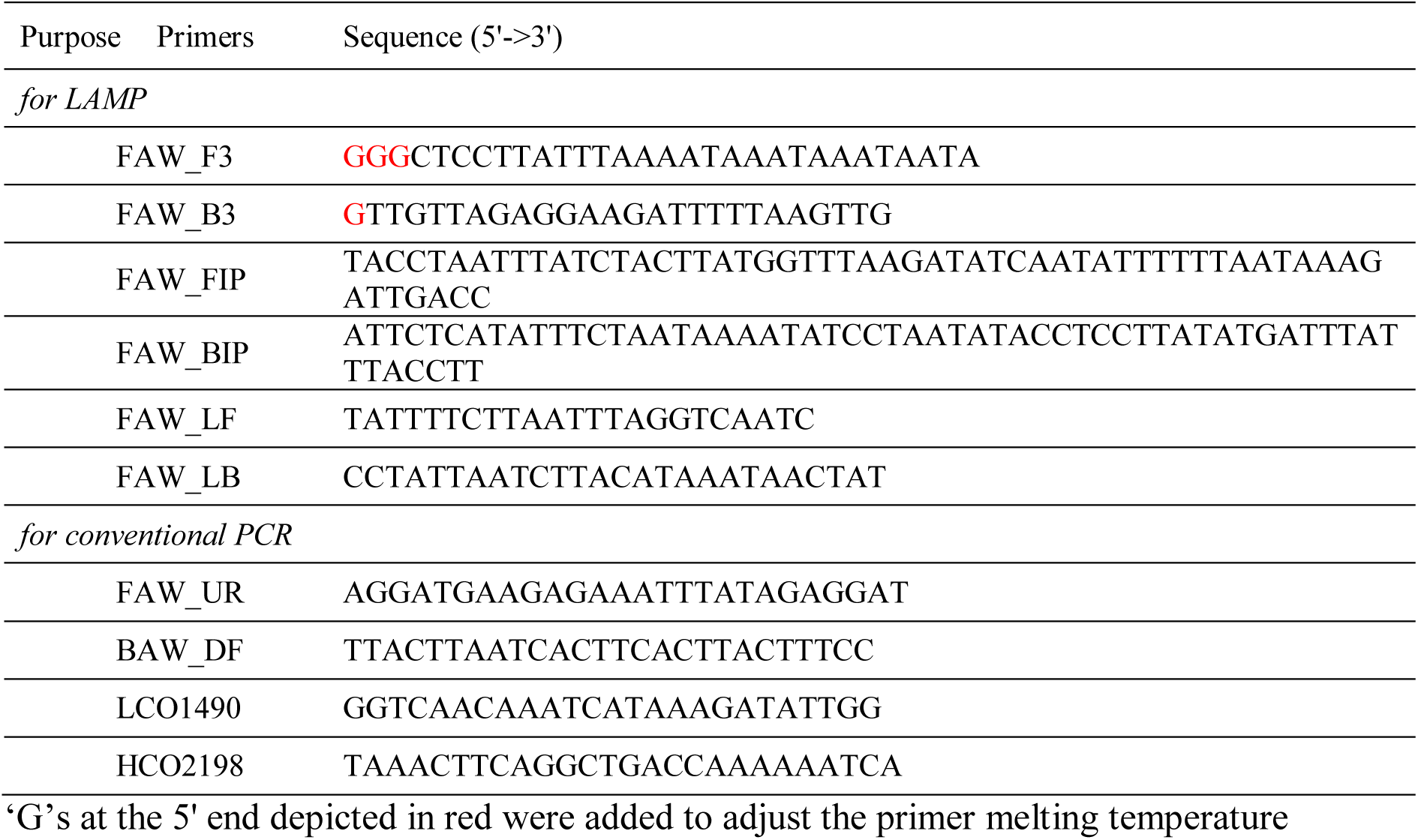
Primer list for LAMP (lamp loop mediated isothermal amplification) and conventional PCR in this study

**Fig. 3.**
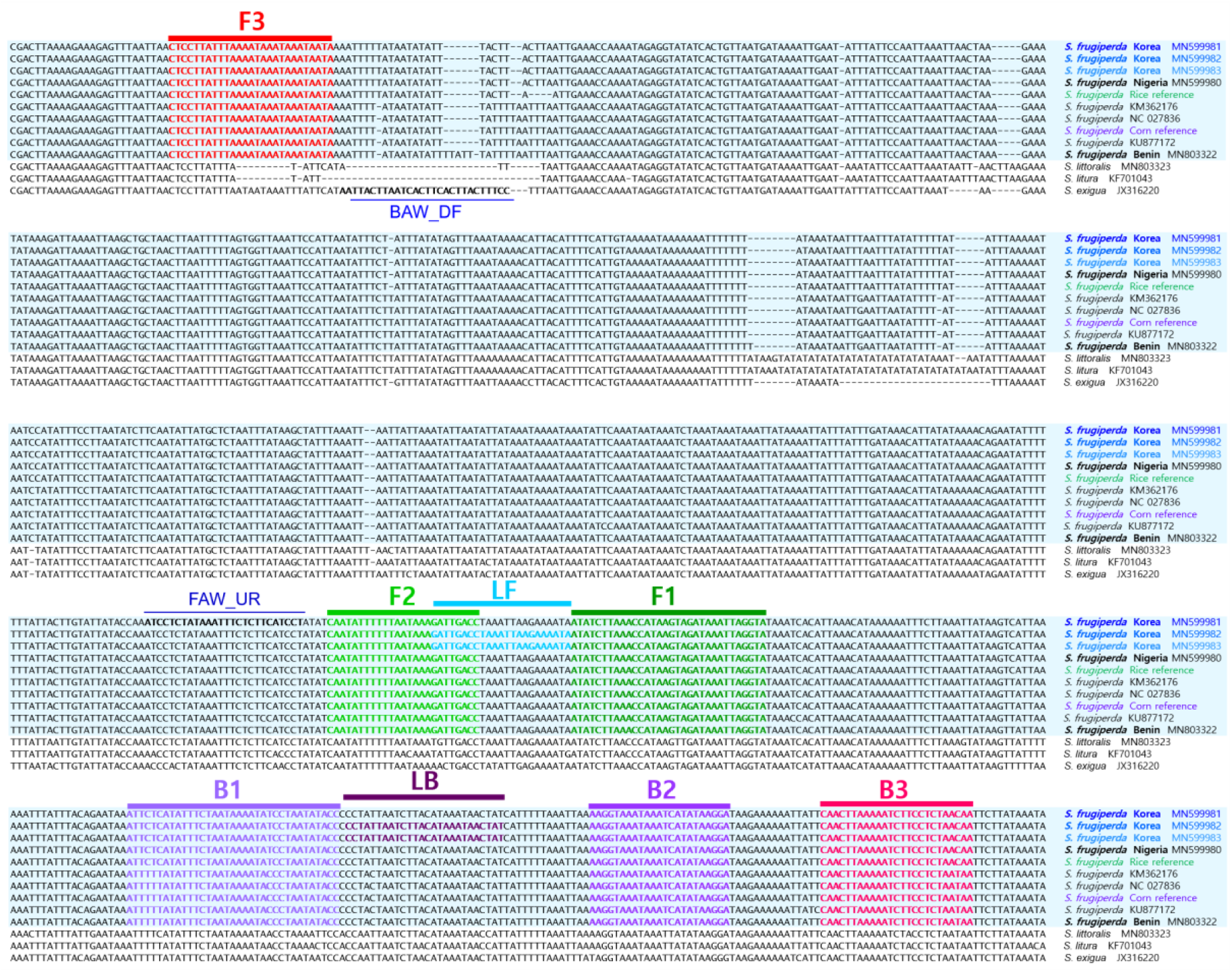
Location of primers and primer binding regions on partial sequence of *Spodoptera* mtDNA. Sequence alignment of several *S. frugiperda* populations *and S. littoralis, S. litura*, and *S. exigua*. Inner primer, FIP is consisted of F1c (complementary sequences of F1) and F2. Another inner primer, BIP is also composed of B1 and B2c (complementary sequences of B2). Essential 4 LAMP primers (F3, FIP, BIP and B3) generate the dumbbell structure and two loop primers, LF and LB accelerate the LAMP reaction (see Nagamine et al. 2002 for details). Used primer information is documented in the Table 2.

**Fig. 4.**
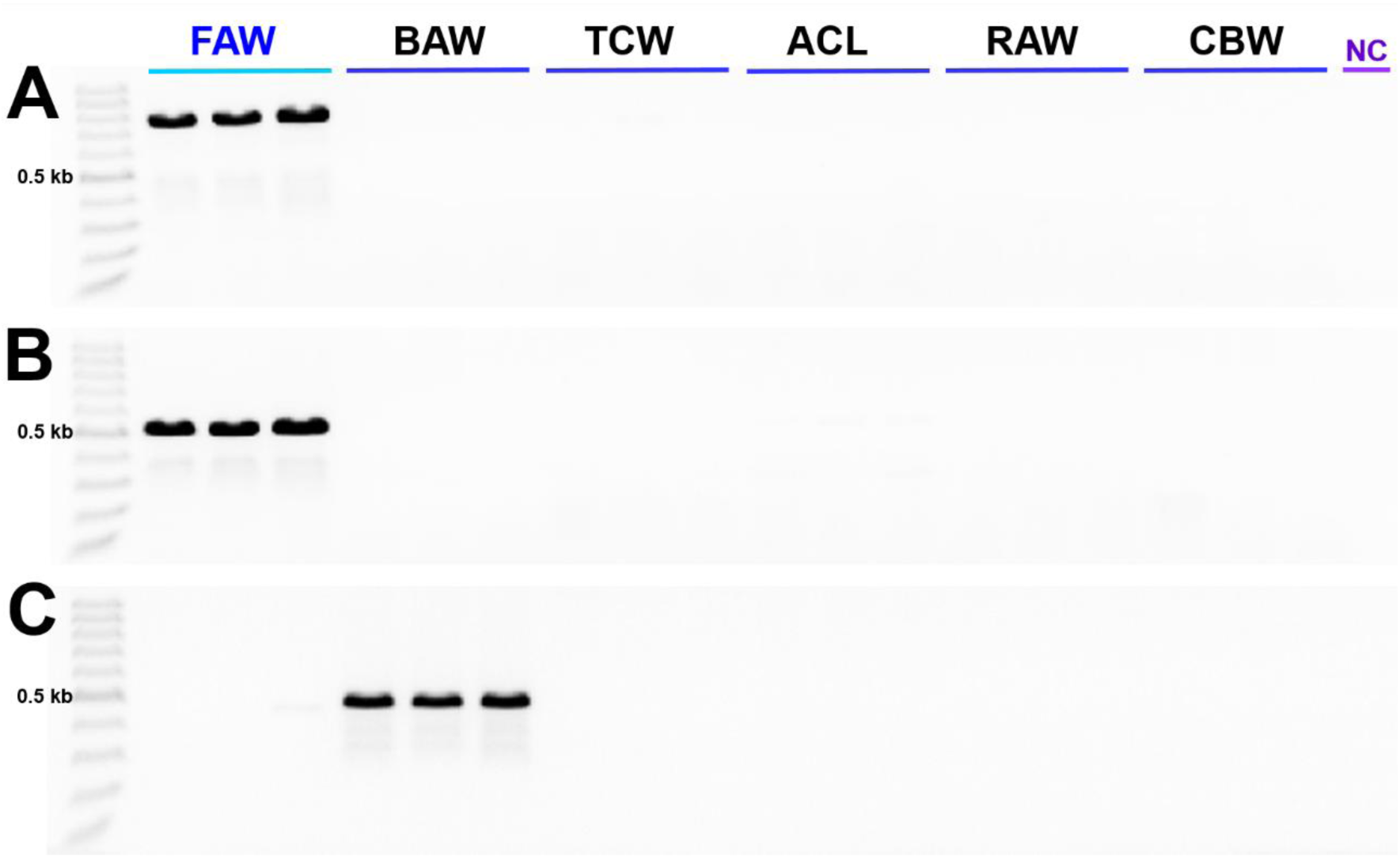
Conventional PCR to distinguish *S. frugiperda* and *S. exigua*. (A) PCR with LAMP external primer set FAW_F3 and FAW_B3 produced a 781 bp product from *S. frugiperda* DNA only. (B) PCR with primer FAW_F3 and alternative reverse primer FAW_UR produed a 501 bp product from *S. frugiperda* DNA only. (C) PCR with primers BAW_DF and FAW_UR produced a 454 bp product from *S. exigua* DNA only. Abbreviations are NC (non-template control), FAW (fall armyworm *S. frugiperda*), BAW (beet armyworm *S. exigua*), TCW (tobacco cutworm *S. litura*), ACL (African cotton leafworm *S. littoralis*), RAW (rice armyworm *Mythimna separata*), and CBW (cotton bollworm *Helicoverpa armigera*).

### 3.2 Conditions for diagnostic PCR and LAMP assay

Standard PCR with diagnostic primers showed that *S. frugiperda* and *S. exigua* could be readily distinguished (Fig. 4). The LAMP assay with four primers was tested with various incubation temperatures ranging from 59 to 65°C. Best results were obtained with 50ng of total DNA, incubated at 61°C for 90 minutes in a 25 μl reaction volume (Fig. 5). A positive reaction can be seen under visible light without any treatment (Fig. 5A) or confirmed under ultraviolet light with Cyber Green (Fig. 5B), or gel electrophoresis (Fig. 5C). Under 100 bp sized bands are not the LAMP product. That’s aggregated primers. So that generated even in negative control. The two loop primers were tested for possible enhancement of the LAMP reaction, as suggested by Nagamine et al. (2002) (Nagamine et al., 2002). Addition of LB enhanced the reaction, producing the same amount of product in ten fewer minutes (Fig. 6C). Addition of LF inhibited the LAMP reaction with LB (Fig. 6D) or without LB (Fig. 6B).

**Fig. 5.**
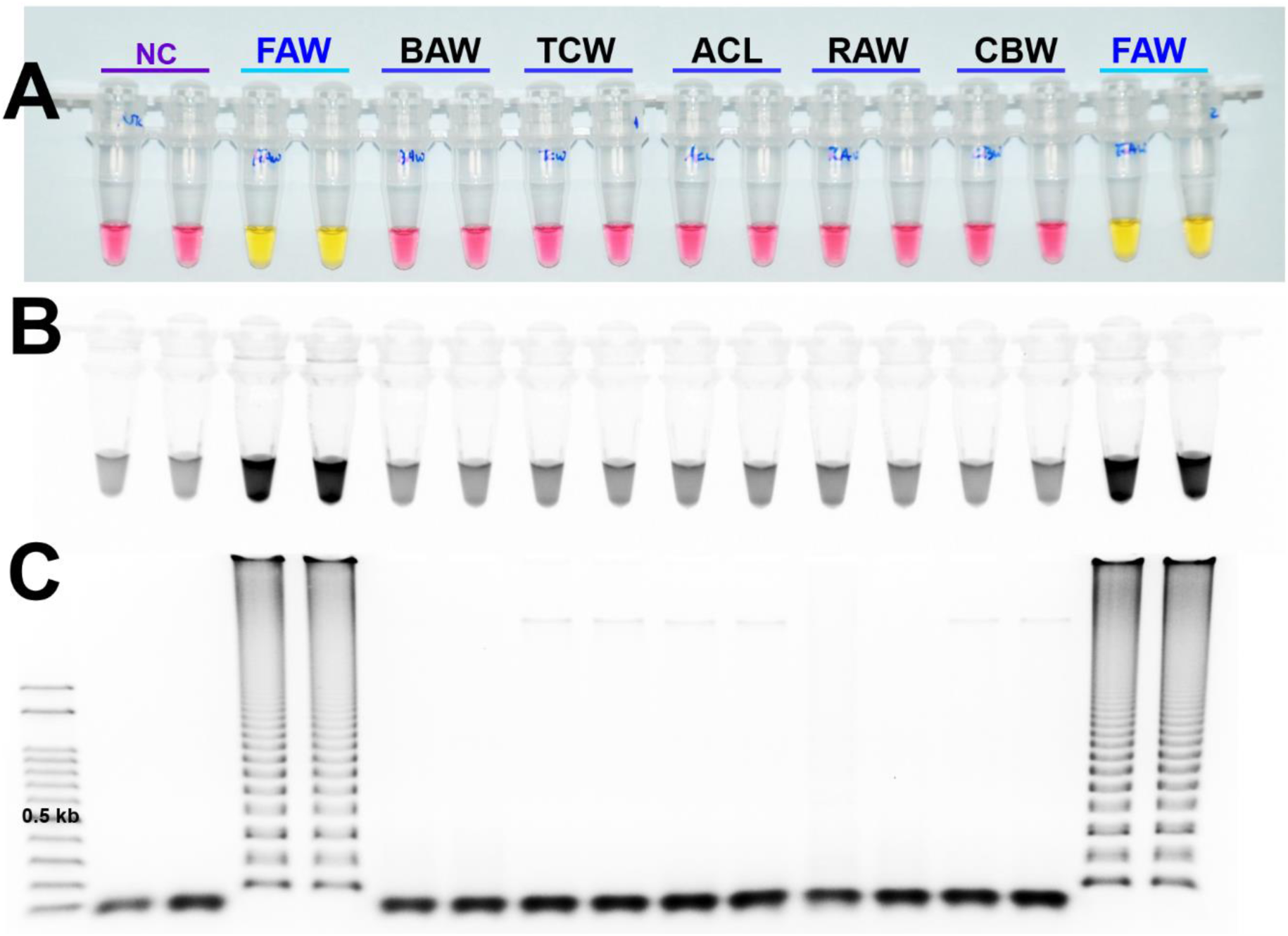
Sensitivity of the LAMP assay results for *S. frugiperda* species detected under (A) visible light, (B) ultraviolet light with Cyber Green and (C) gel electrophoresis. Abbreviations as in Fig. 4. The original pink color of the reaction mixture turned yellow in a positive reaction when product was formed but remained pink in negative reactions (A).

**Fig. 6.**
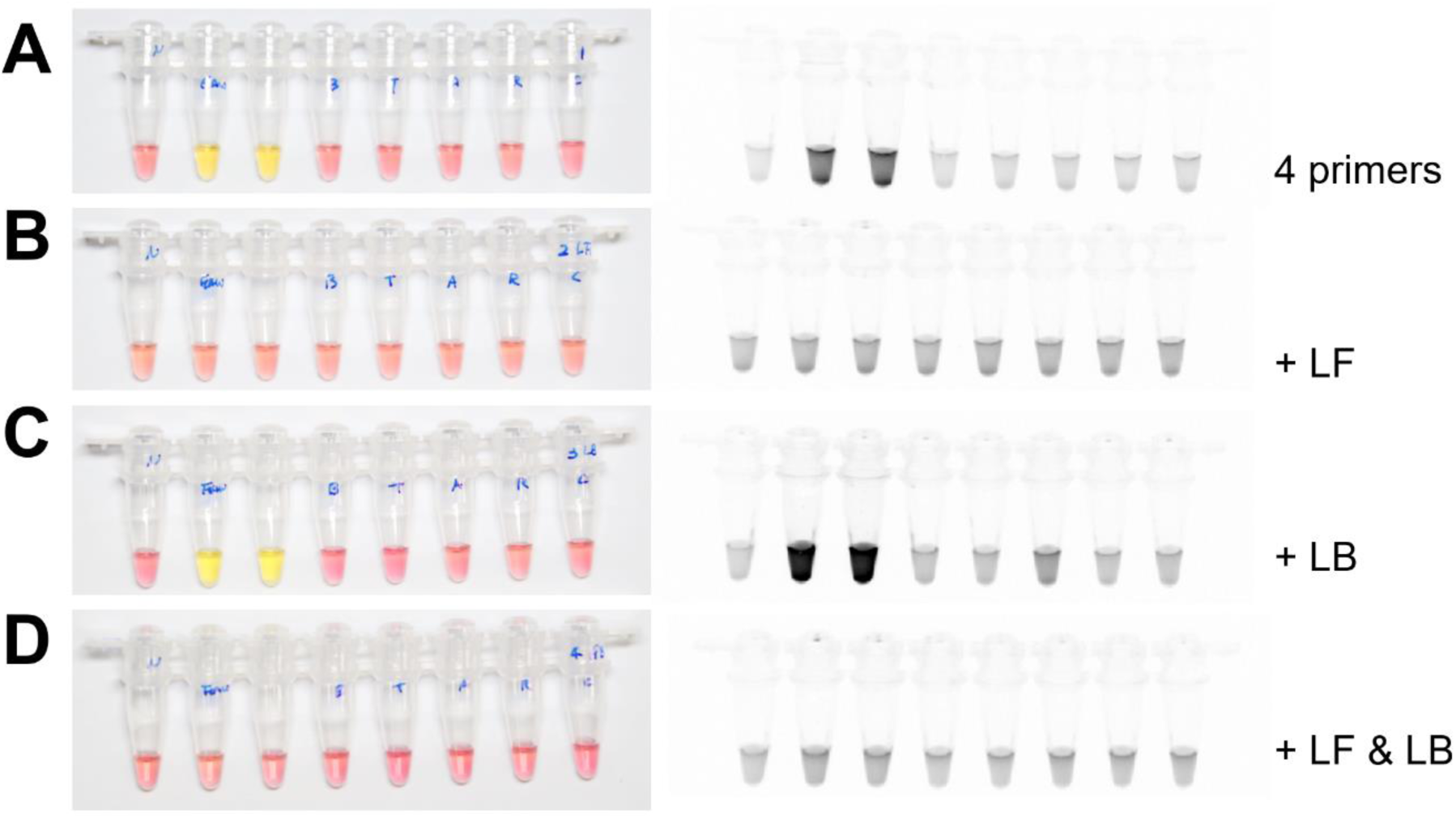
The LAMP assay results with 4 primers and additional loop primers, loop forward, LF and loop backward, LB under visible light (left side) or ultraviolet light with Cyber Green (right side).

The detection limits of the four LAMP primers and LB were tested on purified DNA. The color change under visible light could be detected with as little as 1 ng of DNA (containing genomic DNA and mitochondrial DNA resulting from a standard DNA isolation protocol, Fig. 7A). With UV light in reaction tubes or gel electrophoresis, as little as 10 pg of DNA could be detected (Fig. 7B and C).

**Fig. 7.**
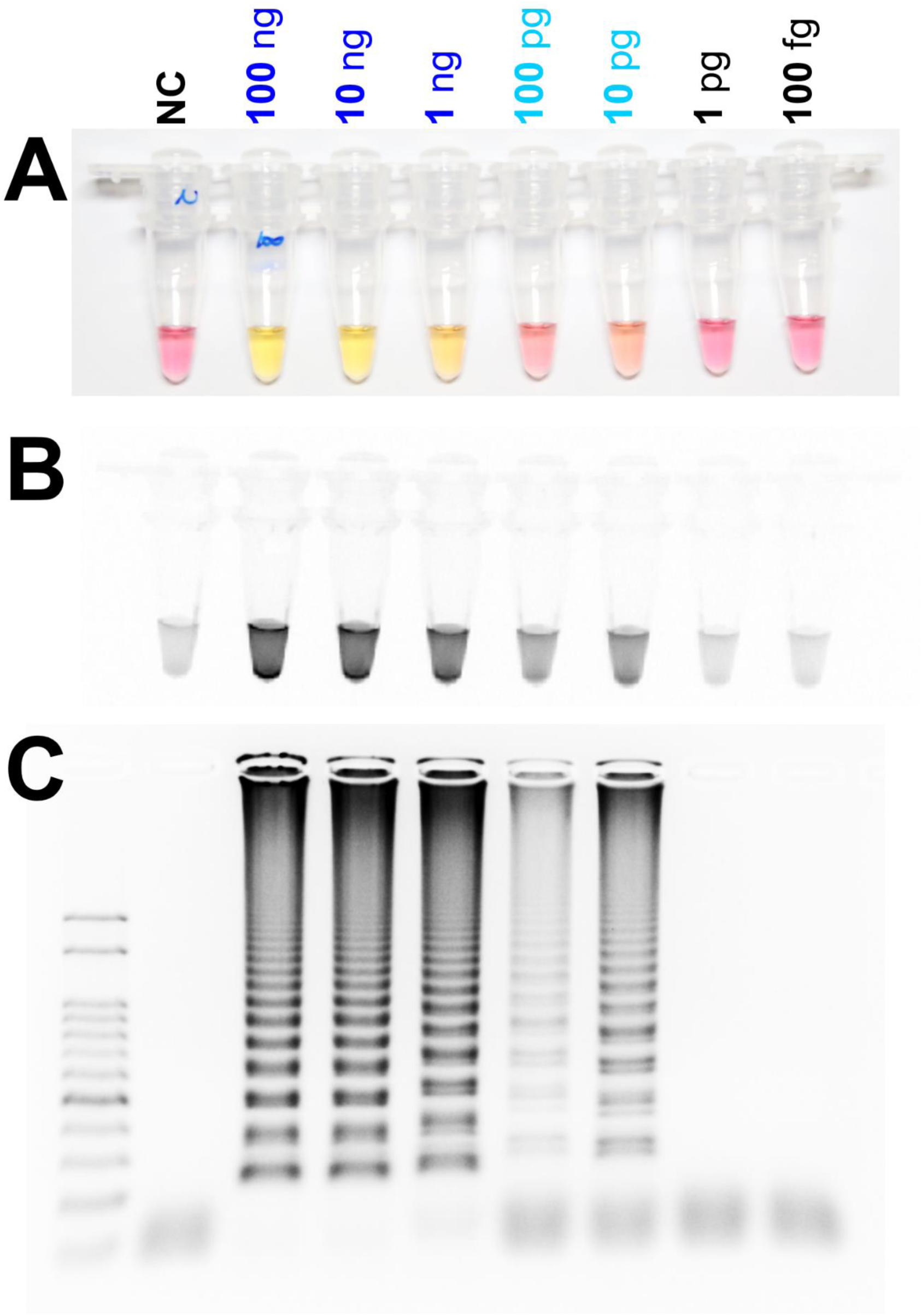
Identification of the detection limit of genomic DNA in the LAMP assay from 100ng to 100fg under (A) visible light, (B) ultraviolet light with Cyber Green and (C) gel electrophoresis.

To test the DNA releasing technique on insect tissue, approximately 10 mg of larval tissue, or an adult leg or antenna, was incubated at 95°C for 5 minutes with 30 μl nuclease free water (Fig. 8B). Then 2 μl of the supernatant was used in the LAMP assay with 5 primers. Results could clearly be seen with visible light (Fig. 8A), UV light with Cyber Green (Fig. 8C) or gel electrophoresis (Fig. 8D).

**Fig. 8.**
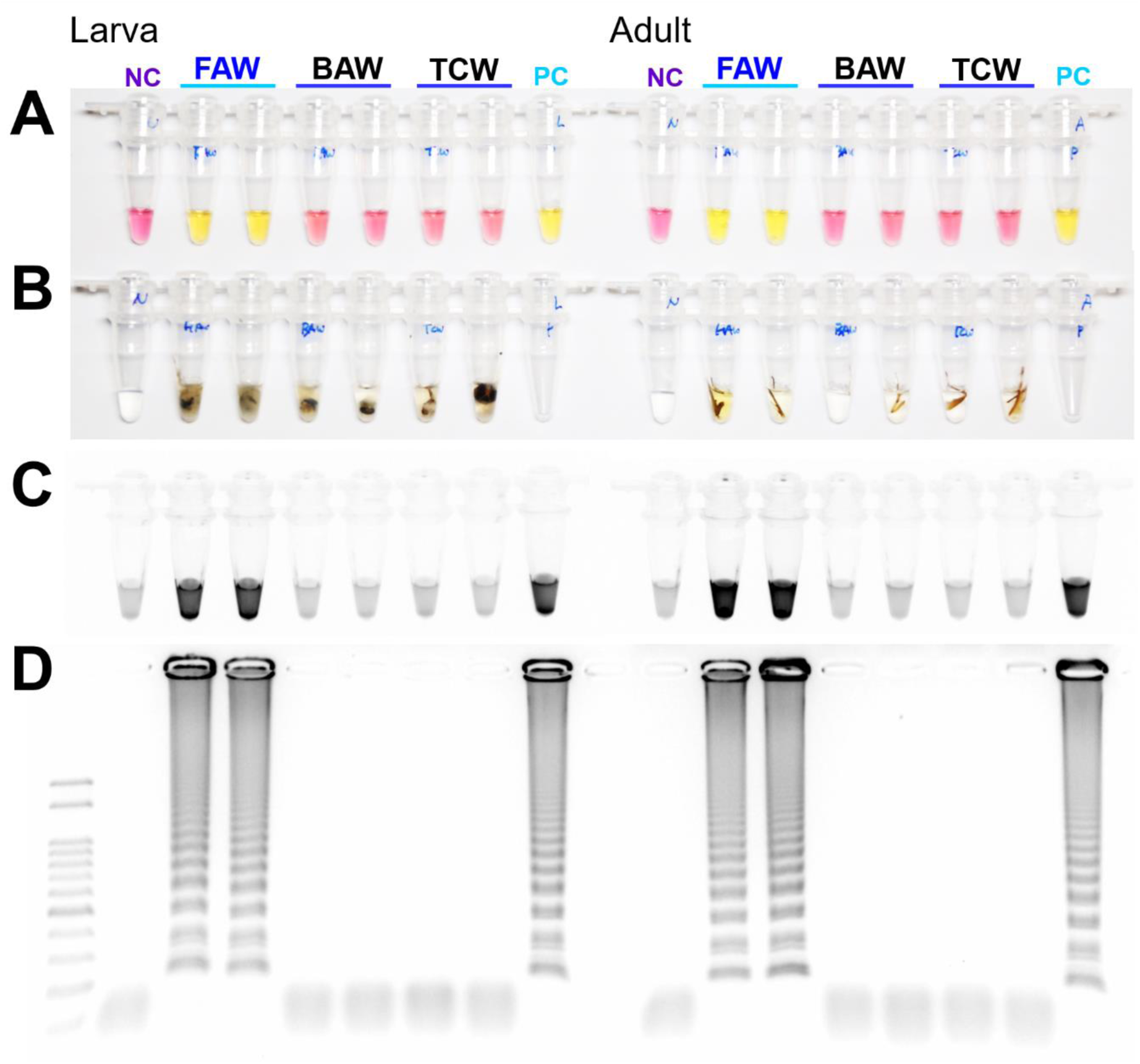
Sensitivity of the LAMP assay results with the DNA releasing technique from (B) insect tissue. (A) LAMP products under visible light, (C) ultraviolet light with Cyber Green and (D) and gel electrophoresis. Around 10mg of larval tissue or adult leg (or antenna) were incubated in 95°C for 5minutes (B). Abbreviations as in Fig. 4, except PC (positive control, isolated DNA from *S. frugiperda*).

## 4. DISSCUSSION

Methods to identify invasive *S. frugiperda* are urgently needed, as farmers and agricultural managers have no prior experience with this pest. The LAMP assay with the modifications described here can fulfil that need. It is not necessary when other species, such as corn borer *Ostrinia* species, can be distinguished morphologically in the field.

As previously reported, the amount of DNA is generally not the limiting factor for the LAMP assay (Choi et al., 2018). Varying the incubation time can generally compensate for different amounts of DNA. The incubation temperature was more important for specificity. At temperatures less than 59°C, *M. separata* produced false positives (data not shown). There are five mismatches between the primer FAW_F3 and the priming sequence of *M. separata* as well as that of *S. exigua*. However, there was no false positive reaction in the optimal incubation condition, 61°C during 90 minutes. On the other hand, incubation temperatures higher than 62°C required longer incubation times to produce the same amount of product (data not shown).

In another attempt to decrease the incubation time, we applied two other loop primers, loop forward (LF) between the F2 and F1 regions and loop backward (LB) between the B1 and B2 regions, which have been suggested to amplify additionally from the loops in the dumbbell forms (Choi et al., 2018). Finding suitable loop primers was difficult because of the high AT content, and the strategy of adding overhanging ‘G’s to the 5’ end to increase the melting temperature was hard to use in the inner primers.

Alternatively, conventional PCR also can be used for *S. frugiperda* species detection as well as *S. exigua* (Fig. 3, Fig. 5). The simplicity, accuracy and adaptability for high throughput of the LAMP assay are distinct advantages (Notomi et al., 2015; Zhang et al., 2014; Mori & Notomi, 2009). Moreover, recently LAMP applied in many ecology studies in and outside of the lab (Lee, 2017).

The origin of *S. frugiperda* populations in Korea has not been identified as of 2019. Based on meteorological predictions, they might migrate from some part of China and East Asia to Korea at least twice (end of May to end of June and July) (Li et al., 2020; Wu et al., 2019). Mitochondrial genome sequencing of three Korean populations implies that there are at least two genetically different populations of possibly different origin which invaded Korea. The next plan of this study is to identify the *S. frugiperda* species using this new LAMP assay and to analyze the identified SNPs, indels and the insecticide resistant related mutations of various field collected *S. frugiperda* populations, which may be useful in tracking gene flow and for insecticide resistance management.

## ACKNOWLEDGEMENTS

The Cooperative Research Program supported this study for Agriculture Science & Technology Development (Project No. PJ01500903), the Rural Development Administration, Republic of Korea. Support was also provided by the Max-Planck-Gesellschaft. Special thanks to Dr. Sabine Haenniger and Dr. Melanie Unbehend in MPI-CE for supplyng the African populations of *Spodoptera frugiperda*. Mr. Hyun Oh Lee in PHYZEN for technical support in NGS and bioinformatics.

## AUTHOR CONTRIBUTIONS

J.K., H.Y.N and M.K. conceived the study. J.K, H.Y.N, M.K., H.J.K., H.J.Y., S.H., M.U. and D.G.H prepared samples and performed mt genome sequencing. J.K., H.Y.N performed experiments. J.K. and D.G.H mainly wrote the paper.

